# A Darwinian Selection Based Explanation of Peacock’s Tail

**DOI:** 10.1101/2020.05.12.086520

**Authors:** Dinkar Wadhwa

## Abstract

There is no satisfactory explanation for why peacock possesses a tail, presence and especially courtship display of which makes the organism vulnerable to predation. Here, I present a model according to which in a polygynous mating system a mechanism which increases vulnerability to predation, a Zahavian handicap, evolves when other two mechanisms to identify high-quality males are either absent or are not sufficiently strong. The two mechanisms are: 1) male resource acquisition ability, and 2) male-male competition for females. The three mechanisms are not necessarily mutually exclusive. Assuming the locus for the tail and choosiness to be sex-specific, it is shown through stochastic simulation that sexual selection, mediated by the tail (a Zahavian handicap), leads to higher rate of increase in the quality of the population of tailed peacocks and tailed-choosy peahens (which exclusively mate with tailed peacocks) as compared to the population of tailless peacocks and tailless-choosy peahens (which exclusively mate with tailless peacocks), through a positive feedback, as daughters of tailed-choosy peahens are of higher average quality and, by virtue of not carrying the tail’s handicap, also fitness than daughters of tailless-choosy peahens. Also, the fold-change in the population of tailed peacocks and tailless-choosy peahens are higher than the fold-change in the population of tailless peacocks and tailless-choosy peahens, for all combinations of the initial conditions. The same results were got, though in milder form, when tailless-choosy peahens were replaced by undiscriminating peahens (which mate with tailless and tailed peacocks in proportion to their frequencies in the population). Although sons of tailed-choosy peahens have lower average fitness than sons of undiscriminating peahens, this difference is inconsequential, because in a polygynous mating system a single male can potentially mate with every female. The work presented here reconciles Zahavi’s handicap principle with Darwin’s theories of natural and sexual selection. Further, it is hypothesized that a genotype responsible for producing in males a reliable indicator of high quality (a Zahavian handicap) or paternal care ability generates mating desirability in females towards males possessing the indicator. It is demonstrated through simulation that this cross-gender pleiotropy expedites the evolution of tailed peacocks and tailed-choosy peahens and leads to higher rate of increase in their quality.

## 1. Introduction

In many species, males possess characteristics such as antlers, ornate tail, and colouration, and behaviours such as courtship display and song which reduce their fitness and thus cannot be explained by the theory of natural selection. In order to explain the presence of such characteristics, Darwin [1] proposed the theory of sexual selection, according to which the lowered fitness of males possessing such characteristics gets offset by their being more attractive to females and thus producing more and/or superiour quality progeny than those males which lack them. But the sexual selection theory did not explain what advantage do progeny of females which choose males possessing such characteristics have over progeny of those females which choose other males, so that the evolution of the female choosiness itself could be accounted for.

According to Fisher [2], initially the male characteristic and male quality were correlated; thus, those females which chose males possessing the characteristic left more descendants than those females which chose males without the characteristic. Thus, in the subsequent generations, males had evolved heightened form of the characteristic and females had evolved higher attraction for it. Then a point arrived when further increase in the characteristic’s size had no correlation with male fitness, but still the increase in the characteristic’s size and female attraction for it continued because of the selective pressure created by the latter. In other words, males possessing the characteristic had more and/or superiour quality progeny simply because of the female attraction for the characteristic. A few questions can be raised on Fisher’s argument: a) in what way the characteristic was correlated with male quality?, b) how that correlation came to an end?, and c) what factor compensates for the cost of the extra size of the characteristic? Additionally, female attraction for a characteristic is not sufficient to cause the evolution and maintenance of the characteristic. This can be seen by considering a hypothetical example where a fraction of females are attracted to, and thus mates with, males which are blue coloured, whereas the other fraction of females mates with colourless males. The blue colour is due to a by-product of defective metabolism. Thus, because of inheriting defective metabolic genes and getting lower quality parental care, the average fitness of progeny of females which mate with blue males is lower than the average fitness of progeny of females which mate with colourless males. This amounts to lower percentage of blue males and females having attraction for blue males in the subsequent generation. Thus, with each passing generation the percentage of blue males and females having attraction for them will decrease.

For a characteristic to undergo positive selection in a polygynous mating system, it should increase fitness of at least female offspring in terms of them getting good genes and/or better paternal care. The characteristic may decrease fitness of male offspring to the extent that only few males survive. The lower survivability of males is inconsequential because in a polygynous mating system a single male can potentially mate with every female. Natural selection can be looked at as a feedback mechanism. Genes that increase fitness (“good genes”) enable the organism to leave more descendants, hence higher presence of the good genes in the subsequent generation. Genes that decrease fitness (“bad genes”) cause the organism to leave fewer descendants, hence lower presence of the bad genes in the subsequent generation. Sexual selection for a characteristic can only escape this loop if the characteristic has no effect on fitness. Thus, Fisher’s argument is tantamount to assuming that the characteristics have no effect on fitness, which is clearly not true.

In 1975, Amotz Zahavi [3] proposed the handicap hypothesis to explain the evolution of the male characteristics and behaviours which, although reduce their fitness, increase their mating success. According to Zahavi, the presence of such fitness-reducing traits are tests, as only high-quality males would be able to survive despite incurring a loss in fitness due to the handicap. Thus, the presence of such traits reliably signals male quality, for males not possessing such traits may or may not be of high quality. It has been demonstrated that offspring of peacocks with more elaborate train grow and survive better under experimental conditions which are approximately the same as natural conditions [4]. In 1976, Maynard Smith [5] analyzed the handicap hypothesis and concluded that the proposed mechanism does not work, even when the handicap is absent in females. In 1978, Bell [6] carried out a more extensive study of the handicap principle, considering haploid population in monogamous and polygynous mating system. The study found out that the principle can work in a certain condition related to the initial frequency of choosy females. The results of Bell’s paper are discussed below. A study by Davis and O’Donald [7] concluded that Zahavi’s model cannot work; but because the analysis in the paper is beyond the scope of the present work, results of that work will not be discussed here.

## 2. Results and Discussion

### 2.1. Fundamental biological functions

In a naive consideration, life can be thought of as an organism’s struggle to leave as many descendants as possible. This, then, demands the organism to succeed in performing at least the following functions: 1) acquire and protect resources, 2) acquire mate, and 3) protect oneself and progeny. For the male gender, the function of mate protection may also be added. In many cases, the female gender has the additional function of gestation to perform. Because resource protection is needed only when resources have been acquired, the function of resource protection may be subsumed into the function of resource acquisition. Further, the functions of protecting progeny and mate (in case mate protection is present) may be subsumed into the need to protect oneself. Arguably, the function of self-protection should also include protecting oneself from parasites and pathogens, because abstractly, parasites and pathogens can be considered equivalent to predators. Also, gestation may be subsumed into resource acquisition. Thus, we are left with three fundamental biological functions, namely resource acquisition, mate acquisition, and self-protection.

In any mating system, female cannot possess characteristics as indicators of quality that reduce their fitness, because reproduction primarily depends on them. In a given species, females which display their quality by a fitness-reducing feature will get outnumbered by females which don’t, because the latter are more likely to survive up to the reproductive age and could invest advantage due to higher fitness in reproduction (higher quality eggs or better gestation) and parental care. In monogamous mating system, because each male can mate with only one female, males also should not indicate their quality by characteristics which reduce fitness. In polygynous mating system, however, because a high-quality male can mate with several females, characteristics that reduce fitness can serve as indicator of quality. This is because a few high-quality males may still survive, and consequently all females can get to mate. Because the present work seeks to explain the evolution of peacock’s tail, the focus will be on how females determine male quality. Thus, a mechanism, or test, needs to exist by means of which females can determine male quality. Naturally, fundamental biological functions of males can serve as the tests. However, because male’s resource acquisition ability may also contribute to his mate acquisition ability, an alternate formulation may be better conceptually. Thus, because the process of mate acquisition consists of a male displaying his ability to acquire resources and/or fighting off rival males, it may be better to replace “resource and mate acquisition” with “resource acquisition and male-male competition”.

a) Resource acquisition: Males may demonstrate their resource acquisition ability in three ways. First, by stockpiling resources. Second, by possessing high-investment characteristics, such as antlers, which are shed and regrown annually. Only those males which are good at resource acquisition will be able to grow and maintain antlers. Third, by building an architecture, e.g. nest or bower. Although vast majority of bird species are monogamous, a bird species may be imagined in which males build multiple nests in order to attract many females. b) Male-male competition: The male which defeats rival males in a physical confrontation demonstrates his high quality. c) Self-protection: This test is implemented by way of males possessing a characteristic or displaying a behaviour which increases their vulnerability to predation. The examples are the peacock’s tail plumes, bright colouration of many bird species, and risky courtship displays and songs. Males which are able to survive in spite of being more vulnerable to predation are of higher quality than those which are not. It should be noted that the presence of increased vulnerability to predation post-mating will hamper paternal care in case it is present. A hypothetical phenomenon can be considered in which males, as part of courtship ritual, put them in a situation which makes them highly vulnerable to parasitic/pathogenic infection. Only high-quality males (i.e., having strong immune system) will be able to survive such ritual. The three functions can also be present in combination. For example, a) antlers are used as weapons in male-male combat for mate acquisition, and b) male-male fighting may also be involved in resource acquisition directly or indirectly, that is, in competition for territorial possession. In the peacock, because the first mechanism is absent and the second (territorial fight) is weakly present, males are tested on their self-protection ability. Therefore, a characteristic, the tail, is evolved in males which makes them more vulnerable to predation. The increased vulnerability to predation exists throughout a peacock’s life, but the model only demands the presence of heightened vulnerability to predation during mate selection process.

### 2.2. The evolution of peacock’s tail and peahen’s choosiness

A haploid population is considered which has i) an autosomal locus for quality, ii) male-specific locus for tail, and iii) female-specific locus which determines the peahen type. The three loci have two alleles. The first locus has Q (whose values are 1.05, 1.1, 1.2, 1.5, 2) and q (whose value is 1). The second locus has T (tailed peacock) and t (tailless peacock). For the third locus two cases are considered. First in which the two alleles are tailed-choosy, peahens which mate only with tailed peacocks, and tailless-choosy, peahens which mate only with tailless peacocks. Second in which the two alleles are tailed-choosy, peahens which mate only with tailed peacocks, and undiscriminating, peahens which mate with both types of peacocks in proportion to their frequencies in the population. In the present model, an additive model of fitness is employed where fitness is equal to quality minus test (the tail’s burden is set equal to q). Because peahens do not have tail, their fitness is equal to their quality, Q. Table 1 shows the equivalence of this model with the additive fitness model of Bell’s paper [6]. The simulation was started with initial population of 10^5^ male and female offspring each. The initial frequency of the allele Q was taken to be 0.01, 0.05, and 0.1. For simplicity, the initial frequency of tailed peacocks and tailed-choosy peahens were considered to be equal, and were taken to be 0.01, 0.05, and 0.1. A tailed peacock having the allele Q reached the reproductive age with the probability (Q-q)/(Q+q), whereas a tailed peacock with the allele q reached the reproductive age with the probability (q-q)/(Q+q); this means that tailed peacocks with the allele q died before being able to reproduce. A tailless peacock or a peahen having the allele Q reached the reproductive age with the probability Q/(Q+q), whereas a tailless peacock or a peahen with the allele q reached the reproductive age with the probability q/(Q+q).

**Table 1:**
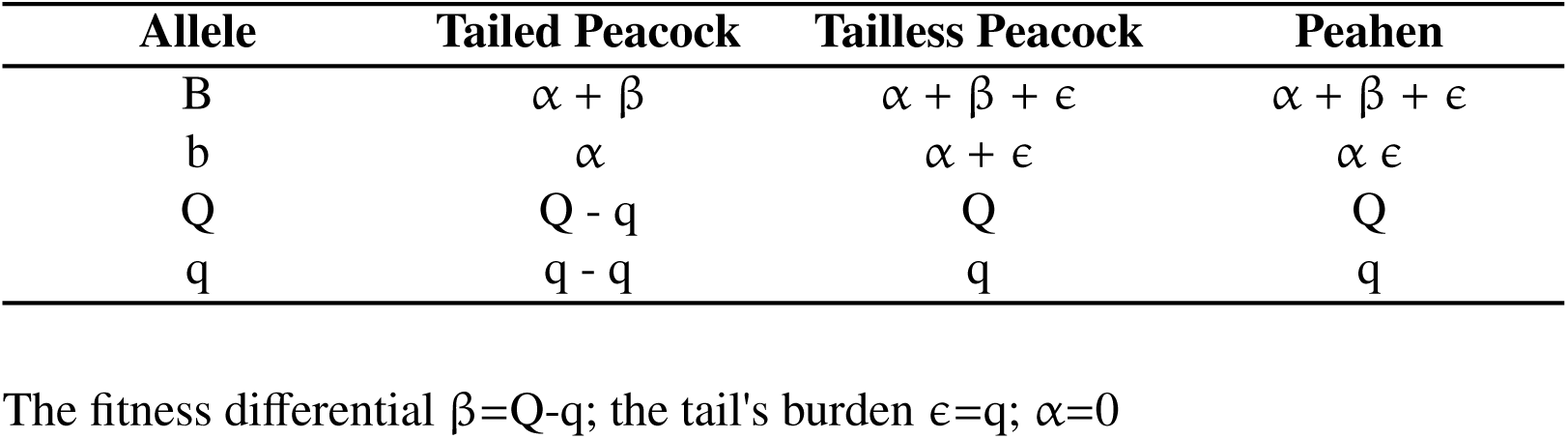
Equivalence of Bell’s additive fitness model [6] and the model in the present work.

For a given value of the fitness differential, the initial frequency of the allele Q, and the initial frequency of tailed peacocks, simulation was carried out for the number of generations at which the value of the below-stated quantities stabilized, because of the fixation of the allele Q by natural selection. The quantities are: 1) the ratio of the fold-change in the population of tailed peacocks to the fold-change in the population of tailless peacocks, 2) the ratio of the fold-change in the population of tailed-choosy peahens to the fold-change in the population of tailless-choosy or undiscriminating peahens, 3) the ratio of the fold-change in the quality of the population of tailed peacocks to the fold-change in the quality of the population of tailless peacocks, and 4) the ratio of the fold-change in the quality of the population of tailed-choosy peahens to the fold-change in the quality of the population of tailless-choosy or undiscriminating peahens. The quality of a population of peafowl is equal to the number of peafowl having the allele Q multiplied by the fitness value of the allele Q plus the number of peafowl having the allele q multiplied by the fitness value of the allele q. In other words, the quality of a peafowl population = (Number of Q peafowls)·Q + (Number of q peafowls)·q.

As argued above, increased vulnerability to predation in peacocks due to the tail enables tailed-choosy peahens to mate with high-quality peacocks, thereby expediting the increase in the quality of the population of tailed peacocks and tailed-choosy peahens. To test this hypothesis, two endogamous populations were considered. The first consisted of tailed peacocks and tailed-choosy peahens, which mate only with tailed peacocks. The second consisted of tailless peacocks and tailless-choosy peahens, which mate only with tailless peacocks. Thus, the second population serves as the control, for natural selection is the only evolutionary force acting upon it. The ratio of the fold-change in the quality of the population of tailed peacocks to the population of tailless peacocks was interpreted as the rate of increase in the quality of peacocks due to sexual selection mediated by the tail. The ratio of the fold-change in the quality of the population of tailed-choosy peahens to the population of tailless-choosy peahens was interpreted as the rate of increase in the quality of peahens due to sexual selection mediated by the tail. The ratio of the fold-change in the population of tailed peacocks to the population of tailless peacocks was interpreted as the rate of the evolution of the tail. The ratio of the fold-change in the population of tailed-choosy peahens to the population of tailless-choosy peahens was interpreted as the rate of the evolution of the female choosiness.

As figure 2 shows, for every combination of the initial conditions all the four fold-change ratios are more than one, demonstrating that the Zahavian handicap leads to higher rate of increase in the quality of peacocks and peahens and expedites the evolution of tailed peacocks and tailed-choosy peahens. The explanation of these results are as follows. The average quality of tailed peacocks which survive to reproduce despite heightened vulnerability to predation is higher than the average quality of tailless peacocks. Therefore, the average fitness of daughters of tailed-choosy peahens is higher than the average fitness of daughters of tailless-choosy peahens. Consequently, more daughters of tailed-choosy peahens survive until the reproductive age as compared to daughters of tailless-choosy, thereby amounting to higher percentage of tailed-choosy peahens in the subsequent generation, and so on. In honour of Amotz Zahavi, I name this positive feedback loop the Zahavian runaway. Alternately, from the perspective of higher survivability of daughters of tailed-choosy peahens, the mechanism proposed here may also be called the fitter-daughter principle. Because these daughters will produce tailed peacocks, the average quality of tailed peacocks also will be higher than the average quality of tailless peacocks, and the percentage of tailed peacocks will also increase with each generation. It should be noted, however, that the peacock percentage is not as crucial a factor as the peahen percentage because the mating system is polygynous, where a single male can potentially mate with every female. The fixation of the allele Q and consequent stabilization of the fold-change ratios does not create a conceptual problem, because the struggle between prey and predator is perpetual. In other words, the prey needs to evolve continuously in order to keep up with the predator’s evolution, and vice-versa. Richard Dawkins reasoned that according to Zahavi’s argument, selection should favour ‘the evolution of males with only one leg and only one eye’ [8]. In the light of the work presented here, the reply is that selection could have favoured the evolution of males with only one leg and only one eye if the genetic mechanism had been such that the defect was phenotypically restricted to males.

**Figure 1:**
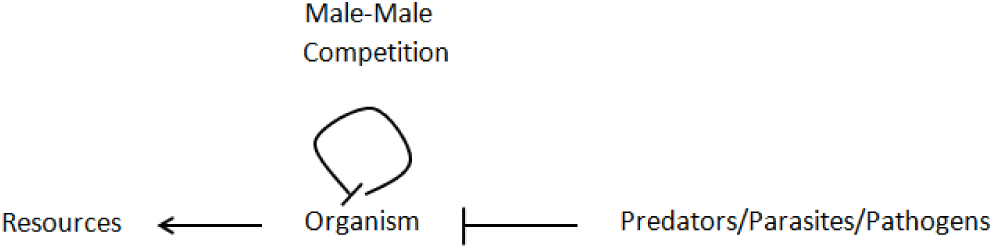
Fundamental biological functions for males in a polygynous mating system.

**Figure 2:**
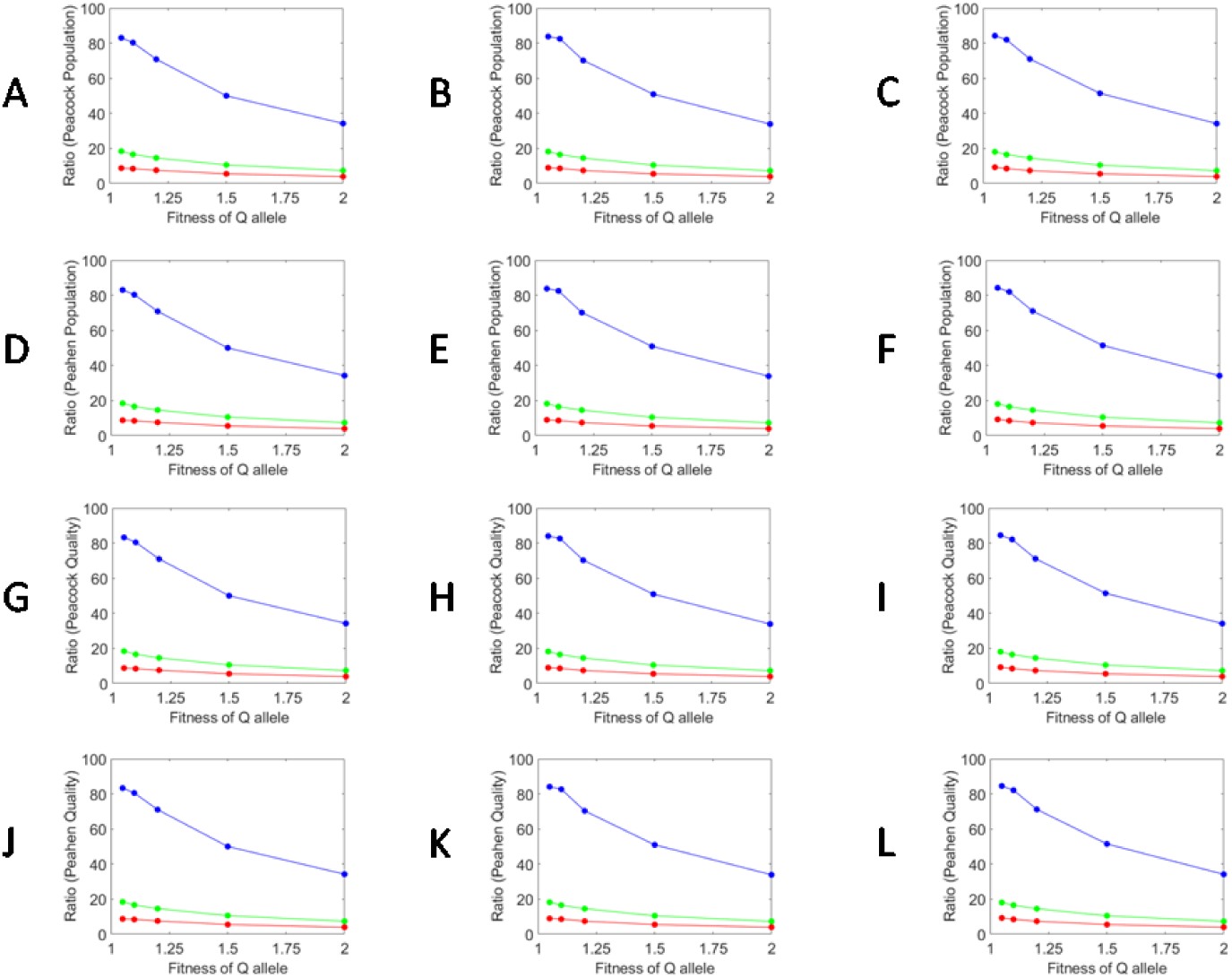
(In the first column, the initial fraction of tailed peacocks and tailed-choosy peahens (f) is equal to 0.01, in the second column f=0.05, and in the third f=0.1. **A**) The ratio of the fold-change in the population of tailed peacocks to the population of tailless peacocks. (**B**) The ratio of the fold-change in the population of tailed-choosy peahens to the population of tailless-choosy peahens. (**C**) The ratio of the fold-change in the quality of the population of tailed peacocks to the quality of the population of tailless peacocks. (**D**) The ratio of the fold-change in the quality of the population of tailed-choosy peahens to the quality of the population of tailless-choosy peahens.

**Figure 3:**
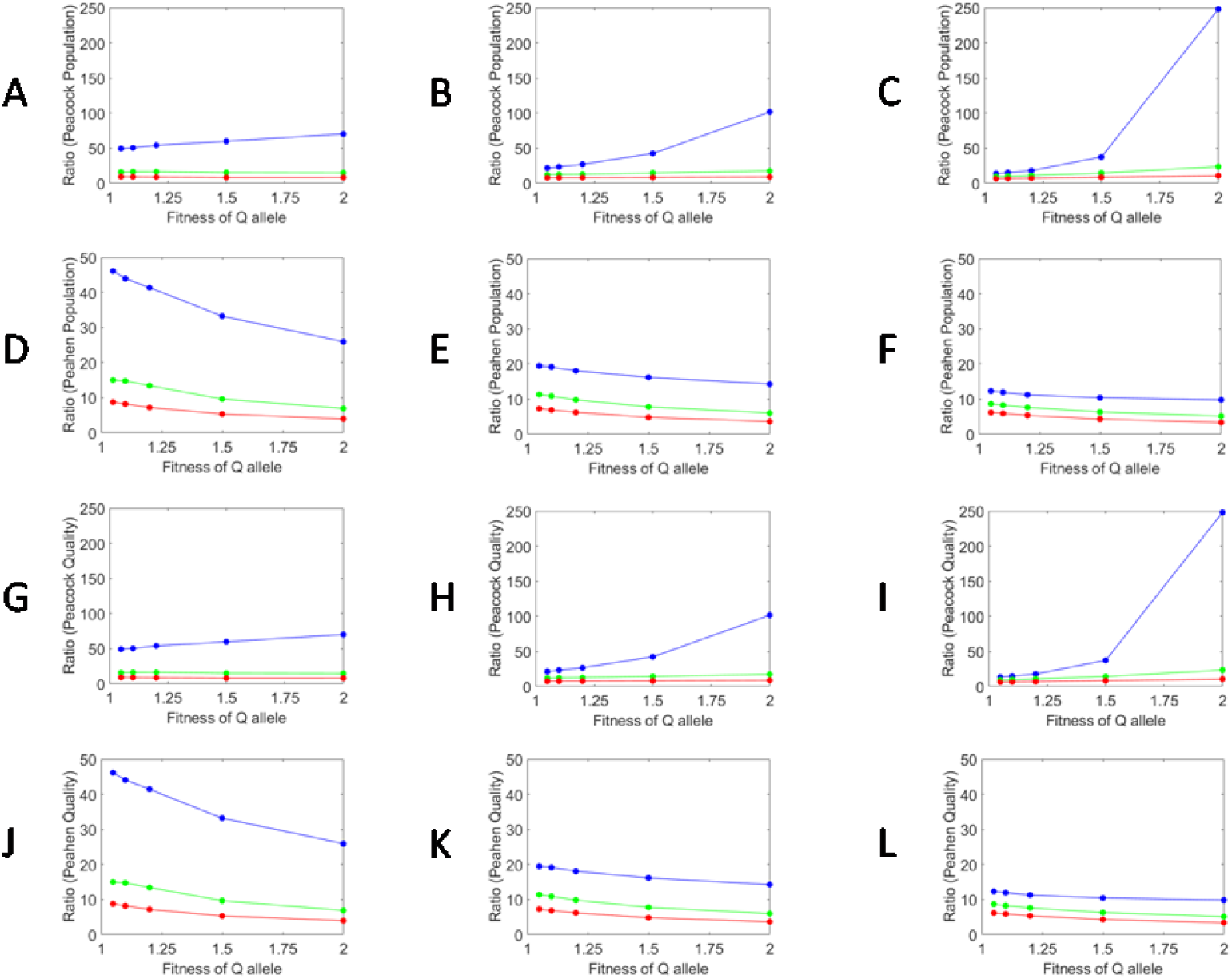
(In the first column, the initial fraction of tailed peacocks and tailed-choosy peahens (f) is equal to 0.01, in the second column f=0.05, and in the third f=0.1. **A**) The ratio of the fold-change in the population of tailed peacocks to the population of tailless peacocks. (**B**) The ratio of the fold-change in the population of tailed-choosy peahens to the population of undiscriminating peahens. (**C**) The ratio of the fold-change in the quality of the population of tailed peacocks to the quality of the population of tailless peacocks. (**D**) The ratio of the fold-change in the quality of the population of tailed-choosy peahens to the quality of the population of undiscriminating peahens.

Notably, all the fold-change ratios decrease as the fitness differential increases. This happens because with higher fitness differential, effect of natural selection becomes stronger (higher percentage of tailless peacocks with the allele Q survives), thereby diminishing the role of the handicap principle, which is mediated through sexual selection. The four fold-change ratios also decrease as the initial frequency of the allele Q increases. The explanation for this trend is as follows. The average quality of tailed peacocks is higher than that of tailless peacocks because tailed peacocks having the allele q do not survive to reproduce, whereas tailless peacocks with the same allele do. With higher initial frequency of the allele Q, fewer tailless peacocks with the allele q are present, thereby reducing the advantage due to the handicap. For very low fitness differentials, the population of tailed peacocks and tailed-choosy peahens went extinct in significant percentages of runs. This was clearly because of very low survivability of tailed peacocks, and thus low chances of mating between tailed peacocks and tailed-choosy peahens, when the fitness differential is low. However, once the two populations sustained up to the sixth generation, they never went extinct.

The same analysis was carried out by replacing tailless-choosy peahens with undiscriminating peahens, which mate with tailed and tailless peacocks in proportion to their frequencies in the population. As previously, for every combination of the initial conditions all the four fold-change ratios are more than one, though there are a few differences. For all combinations of the initial conditions, the ratio of the fold-change in the population of tailed-choosy peahens to the population of undiscriminating peahens is lower than the ratio of the fold-change in the population of tailed-choosy peahens to the population of tailless-choosy peahens. The explanation for this is as follows. Because the average quality of tailed peacocks is higher than the average quality of tailless peacocks, the average quality (and hence also average fitness) of daughters of undiscriminating peahens is higher than the average quality of daughters of tailless-choosy peahens. Consequently, more daughters of undiscriminating peahens survive to reproduce, which lowers the fold-change ratio. Further, as the initial fraction of tailed peacocks increases, the ratio of the fold-change in the population of tailed-choosy peahens to the population of tailless-choosy peahens remains constant, but the ratio of the fold-change in the population of tailed-choosy peahens to the population of undiscriminating peahens decreases. This is because the likelihood of mating between undiscriminating peahens and tailed peacocks goes up as the initial fraction of tailed peacocks increases.

For low fitness differentials, the ratio of the fold-change in the population of tailed peacocks to the population of tailless peacocks is lower for the case where undiscriminating peahens are present as compared to the case where tailless-choosy peahens are present. For higher values of the fitness differential, the reverse is true. Specifically, in the undiscriminating peahens case, although the number of tailed peacocks is higher, the increase in the number of tailless peacocks is such that the ratio of the fold-change in the population of tailed peacocks to the population of tailless peacocks is lower than the tailless-choosy peahens case, for low fitness differentials. The reason for this result is as follows. Mating between tailed peacocks and undiscriminating peahens has two opposing effects on the growth rate of tailless peacocks: 1) it reduces the growth rate of tailless peacocks by lowering their chances of mating with undiscriminating peahens, and 2) it increases the growth rate of tailless peacocks in the following way. Because the average quality of tailed peacocks is higher than the average quality of tailless peacocks, mating of undiscriminating peahens with tailed peacocks, instead of tailless peacocks, produces higher quality undiscriminating peahens. In the next generation, mating between tailless peacocks and undiscriminating peahens which received the higher quality allele, Q, from their tailed father produces higher quality tailless peacocks. When the fitness differential is high, the former effect dominates because the quality of tailless peacocks increases much more due to natural selection, whose intensity is higher due to large fitness differential, than due to the coupling between tailless peacocks and tailed peacocks via undiscriminating peahens, and vice-versa. The reasoning also explains the result that although the fold-change in the population of undiscriminating peahens increases for all fitness differentials as the initial fraction of tailed peacocks goes up, the fold-change in the population of tailless peacocks, as the initial fraction of tailed peacocks goes up, decreases only when the fitness differential is equal to 2.

However, for all fitness differentials, the fold-change in the population of tailed peacocks does not increase as the initial fraction of tailed peacock goes up. This was unexpected because as the initial fraction of tailed peacock increases, more mating events occur between tailed peacocks and undiscriminating peahens. It seems that this discrepancy might be due to very high variance of the fold-change in the population of tailed peacocks at low initial frequency of tailed peacocks. The variance decreased as the initial frequency of tailed peacocks increased. For the initial frequency of tailed peacocks equal to 0.01, the variance and mean had the same order of magnitude, whereas for the initial frequency equal to 0.1, the variance was one order of magnitude less than the mean. The same trend was observed for the fold-change in the population of tailed peacocks for the tailless-choosy peahens case and also for tailed-choosy peahens in both cases (i.e., tailless-choosy peahens and undiscriminating peahens cases). In any case, because the ratio of the fold-change in the population of tailed-choosy peahens to the population of undiscriminating peahens is more than one, the evolution in this mating scenario also occurs through Zahavian runaway.

The study by Bell [6] showed that the handicap principle can work within a certain range of the initial frequency of choosy females. Specifically, for a given value of the fitness differential (which is equal to Q-q in the present work), there will be a value of the initial frequency of tailed-choosy females, below which the handicap principle cannot not work. However, according to the mechanism proposed here, whatever may the initial frequency of tailed-choosy females be, the handicap principle will always work. That’s because the average fitness of daughters of tailed-choosy females will always be greater than the average fitness of daughters of undiscriminating females. This leads to higher percentage of tailed-choosy peahens, and therefore lower percentage of undiscriminating peahens, in the subsequent generation.

A study [9] showed that peacocks which exhibited frequent behavioural display and had a large number of tail eyespots had better immune system. Peacocks which displayed frequently put themselves at predation risk more often than those which displayed less frequently. Arguably, this behaviour should be heritable. A tail eyespot most probably gives a semblance of a face, and consequently invites more attention from predators. Thus, frequent behavioural display and more tail eyespots translate to stronger test. Only high-quality peacocks are able to pass this test, and daughters of those peahens which mate with such peacocks have higher fitness. Overtime, this will lead to a correlation between high quality (such as better immune system) and frequent behavioural display and more tail eyespots. This result may seem to support the Hamilton-Zuk hypothesis, but an alternative, more general explanation is provided below.

### 2.3. The Hamilton-Zuk hypothesis

According to the Hamilton-Zuk hypothesis, male secondary sexual characteristics signal resistance to parasites and pathogens [10]. Recently, a comparative genome analysis of Indian peacock [11] reported that the immune system genes of the bird displayed signs of adaptive evolution. The authors observed this result of being supportive of the Hamilton-Zuk hypothesis. However, a more general reasoning can account for the evolution of the immune system genes in peacock [11] and the correlation reported between better immune system and higher frequency of behavioural display and large number of tail eyespots [9], if infection by parasites or pathogens are considered to be a test. A peacock with weak immune system and carrying an infection will sluggishly respond to a predatory attack and consequently will be less likely to survive. This will lead to selection of higher quality immune system. Further, all faculties involved in evading predation, such as the sensory systems and muscles and bones involved in taking wing and flying, will undergo selection for improvement. This may explain the findings of the comparative genome analysis [11] that genes involved in bone morphogenesis and skeletal muscle development showed signs of adaptive evolution.

### 2.4. Cross-gender pleiotropy

The analysis above is based on the assumption that the locus for the tail is male-specific and the locus which determines peahen type (tailed-choosy, tailless-choosy, and undiscriminating) is female-specific. However, because an ovariectomized peahen develops peacock-type plumage [12], the locus for the tail has to be autosomal. If the tail’s locus is assumed to be autosomal, half of the daughters (and also sons) of tailed-choosy peahens will be having the T allele. Therefore with each passing generation the percentage of tailed-choosy peahens having the t allele will get halved. Thus it is not clear why the tail’s locus should be autosomal. This led the author to consider a model where all three loci were autosomal and peahens were either tailed-choosy or undiscriminating. Although, due to the lack of computing power, the simulation could not be done for the number of generations at which the four fold-change ratios stabilize, it could still be observed that the ratios were very low and most likely were not going to become more than one. Mechanistic basis for this result is straightforward: because the initial population of tailed peacocks (and tailless peacocks) had the undiscriminating allele; initially, 50% of the daughters from mating between a tailed peacock and tailed-choosy peahen were undiscriminating.

As a result, the advantage that tailed-choosy peahens had over undiscriminating peahens in terms of mating exclusively with high-quality peacocks was no more present. 25% of the sons from such a mating will be tailless peacocks having the tailed-choosy allele and another 25% will be tailed peacocks having the tailed-choosy allele. Although half of the daughters from mating between peacocks having the tailed-choosy allele and an undiscriminating peahen will be tailed-choosy, such mating will be very rare to contribute to the evolution of tailed-choosy peahens. The negative results got when the loci for the tail and choosiness are considered to be autosomal leads us to the question: what could be a purpose of the tail’s locus being autosomal when normal peahens do not have the tail? I hypothesize that genotype for the tail in peahens contributes, partially or fully, to their desirability to mate with a tailed peacock. Generalizing, a genotype responsible for producing a Zahavian handicap in males generates mating desirability in females towards males possessing the Zahavian handicap. Because, besides good genes, parental care can also increase fitness of the progeny, the hypothesis can be extended. A genotype responsible for producing in males a reliable indicator of high quality (a Zahavian handicap) or paternal care ability generates mating desirability in females towards males possessing the indicator.

In the earlier simulation, all daughters from a mating between a tailed peacock and an undiscriminating peahen were undiscriminating. By contrast, when genotypes for the tail and female choosiness are assumed to be the same, half of the daughters from mating between a tailed peacock and an undiscriminating peahen are tailed-choosy. Although, half of the sons also are tailless, but that is immaterial in a polygynous mating system. Thus, when genotype for the tail and female choosiness are identical, the evolution of tailed peacocks and tailed-choosy peahens should be faster. Also, the rate of increase in the quality of tailed peacock and tailed-choosy peahens should be higher. If these predictions are correct, then the four fold-change ratios of the case where genotype for the tail and female choosiness are identical should be higher than the respective four fold-change ratios of the case where one fraction of peahens are tailed-choosy while another fraction is undiscriminating, for all combinations of the initial conditions. Therefore, the ratio of the two ratios should be greater than one. The simulation, however, could not be carried out for the number of generations at which the four fold-change ratios stabilize, because of the lack of computing power. But even for lower number of generations, the prediction turns out to be true, though the result are not presentable.

## 3. Conclusion

There are at least two reasons why Zahavi’s handicap principle found it hard to gain acceptance. Firstly, Zahavi did not state the conditions in which males of a species evolve a handicap, and secondly, he proposed that the evolution of handicaps occur through a non-Darwinian mechanism [13]. Zahavi later suggested that high-quality males may incur lower reduction in fitness than low quality males for the same sized handicap [14]. In simulations done so far, tailed peacocks with the allele Q survived up to the reproductive age with the probability (Q-q)/(Q+q), whereas the survival probability for tailless peacocks was zero (i.e., (q-q)/(Q+q)). In other words, the fitness loss was equal for high- and low-quality males. The extreme of Zahavi’s suggestion would be to assume that the handicap causes no fitness loss in high-quality males. Thus simulations were done assuming that the handicap in males with the allele Q did not cause a loss in fitness (i.e., the survival probability equal to Q/(Q+q)). The results got were the same as those got when the fitness loss was equal for high-quality males and low quality males (data not shown). This was because, as has been mentioned above, in a polygynous mating system only one male needs to survive in order for all females to get the opportunity to mate. Hence it is shown that a handicap need not cause lower reduction in fitness of high-quality males to make the handicap principle more likely to work.

Additionally, the work presented here also hypothesizes that genotype responsible for producing a handicap in males generates mating desirability in females to mate with males possessing the handicap. This hypothesis has an important implication in the evolution of Zahavian handicaps. Simulations carried out in this work assume the presence of an initial population of peahens which exclusively mate with tailed peacocks, besides the presence of an initial population of tailed peacocks. The need for the simultaneous presence of males possessing a Zahavian handicap and females having mating desirability towards such males makes the evolution of a male handicap and female attraction towards it less likely than if genotype responsible for producing the handicap in males and generating attraction towards it in females is assumed to be identical. In nature, however, genotypes for the two traits need not be identical to produce the evolutionary advantage. Even if genotype in male responsible for producing the handicap generates the mating desirability in females with a moderate probability (say, 0.5), the evolution of the handicap and female attraction towards it will still be faster than when the cross-gender pleiotropy was absent.

## Notes

### Competing Interest Statement

The authors have declared no competing interest.

